# Coagulation factor XII contributes to renin activation, heart failure progression, and mortality

**DOI:** 10.1101/2024.09.06.611753

**Authors:** Inna P. Gladysheva, Ryan D. Sullivan, Sofiyan Saleem, Francis J. Castellino, Victoria A. Ploplis, Guy L. Reed

**Affiliations:** Department of Internal Medicine and Translational Cardiovascular Research Center, University of Arizona College of Medicine–Phoenix, Phoenix, AZ, USA; Department of Chemistry and Biochemistry and W. M. Keck Center for Transgene Research, University of Notre Dame, Notre Dame, IN, USA

## Abstract

Symptomatic heart failure (sHF) with cardiac dysfunction, edema, and mortality are driven by overactivation of the renin-angiotensin-aldosterone system (RAAS). Renin is widely recognized as a key initiator of RAAS function, yet the mechanisms that activate renin remain a mystery. We discovered that activated coagulation factor XII generates active renin in the circulation and is directly linked to pathological activation of the systemic RAAS, development of sHF, and increased mortality. These findings suggest a new paradigm for therapeutically modulating the RAAS in sHF and other pathological conditions.

---

Despite improvements in treatment, heart failure with reduced ejection fraction (HF) remains a frequent cause of hospitalization, disability, and death worldwide.^1^ In HF, there is pathologic overactivation of the systemic renin-angiotensin-aldosterone system (RAAS), which harms the heart, promotes symptomatic edema, and contributes to mortality.^2-4^ RAAS activation in the circulation is triggered by the enzyme renin.^3-6^ Levels of plasma renin activity are elevated in patients and mice with dilated cardiomyopathy (DCM) with symptomatic HF (sHF) without chronic renal dysfunction.^4,7-11^ Although renin was discovered more than a century ago, and acknowledged as a master regulator of these processes, the mechanism(s) that generate the active enzyme renin, from its zymogen in vivo, remains a mystery (**Figure 1**).

**Figure 1.**
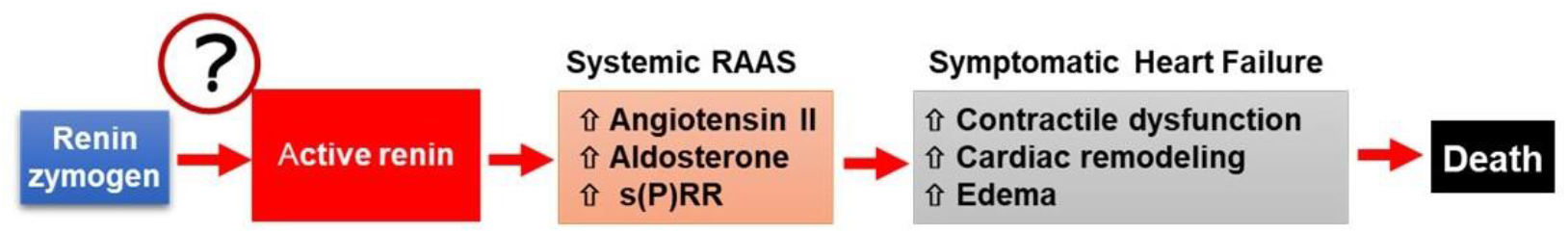
Schema shows that active renin or renin plasma activity is a master regulator of the activation of the systemic RAAS and the progression of HF with reduced EF.

In vitro, renin can be activated from the renin zymogen (pro-renin) by several proteases, including active factor XII (FXII). ^6,12,13^ We examined whether levels of FXII were altered in DCM patients with sHF. FXII levels were increased and positively correlated with pathologically increased levels of plasma renin activity^7^ (**Figure 2**, Spearman *r* = 0.37, P* = 0.013).

**Figure 2.**
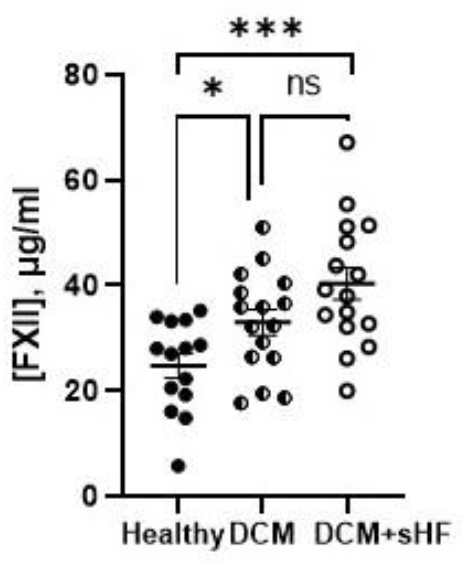
FXII immunoreactive levels are significantly increased in the plasma of a prospective patient cohort with symptomatic DCM and chronic HF without kidney dysfunction compared to levels in healthy patients. Patients’ groups: healthy – normal EF and no HF; DCM (dilated cardiomyopathy) without HF symptoms; DCM + sHF (HF symptoms). Data were analyzed with unpaired Mann-Whitney test using GraphPad Prism 9.5 software and presented as the mean ± SE, * P < 0.5, ***P<0.01. Patients’ enrollment and detailed inclusion/exclusion criteria were previously described.^7,20^

Investigating generation of renin activity in circulation in vivo has been difficult because renin activity levels are low and minimally modulated under physiological conditions. To overcome these limitations, we used a well-established model of DCM that simulates progressive human HF, in which mice transition from “at-risk” to “symptomatic” HF with significantly elevated renin activity plasma levels.^4,7,9-11,14,15^ To determine the potential role of active FXII in generating renin activity in circulation, we backcrossed DCM and FXIII-deficient (FXII-/-)^16,17^ mice (both on a C57BL/6J background) for several generations. Genetically-verified offspring on the DCM background were enrolled in studies at 4 weeks of age, when the mice were “at risk” for DCM but still had normal heart function. Mice were reanalyzed at 13 weeks when the DCM mice normally progress to sHF.^7,9,11,15,18^ We demonstrated in vitro that exogenous active FXII generates renin activity in the plasma of DCM mice with FXII-deficiency and that inhibition of FXII activity blocks this activation (**Figure 3 A-B**). We confirmed in vivo that plasma renin activity levels were robustly reduced in the DCM mice with FXII-deficiency versus the DCM control group (**Figure 4 A**), and plasma levels of immunoreactive, total renin (i.e., predominantly renin zymogen) were not modified (**Figure 4 B**).

**Figure 3.**
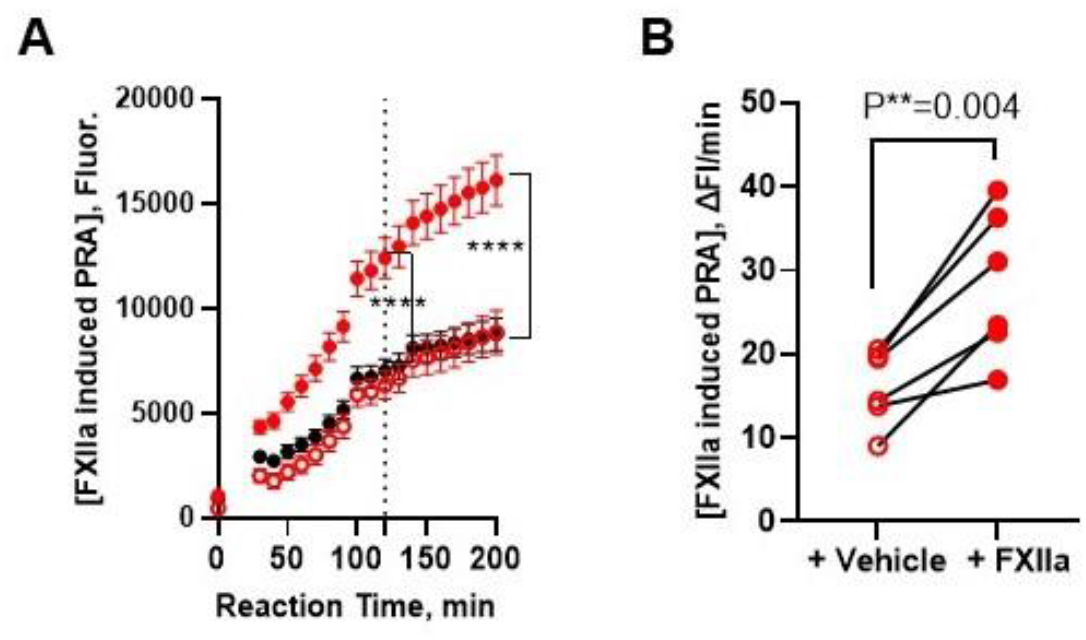
Generation of renin activity by exogenous active FXII in DCM mouse plasma with FXII-deficiency in vitro. (**A**) Exogenous active FXII (FXIIa) generates renin activity (PRA) in the plasma of DCM mice with FXII-deficiency in the presence of vehicle (red closed circle) or a specific inhibitor of FXIIa, H-D-Pro-Phe-Arg-chloromethylketone (black closed circle) that irreversibly inhibits the enzymatic activity of FXIIa versus control plasma (vehicle without FXIIa/inhibitor, red open circle). Data (triplicates) show plasma renin activity (PRA) generation over time in pooled plasma (n = 6). (**B**) Exogenous FXIIa generates PRA in the paired plasma samples of DCM mice with FXII-deficiency (n = 6 mice) vs vehicle. Data were analyzed with unpaired (**A**) or paired (**B**) Mann-Whitney test using GraphPad Prism 9.5 software and presented as the mean of reaction velocity ± SE. **P<0.01, ****<0.0001.

**Figure 4.**
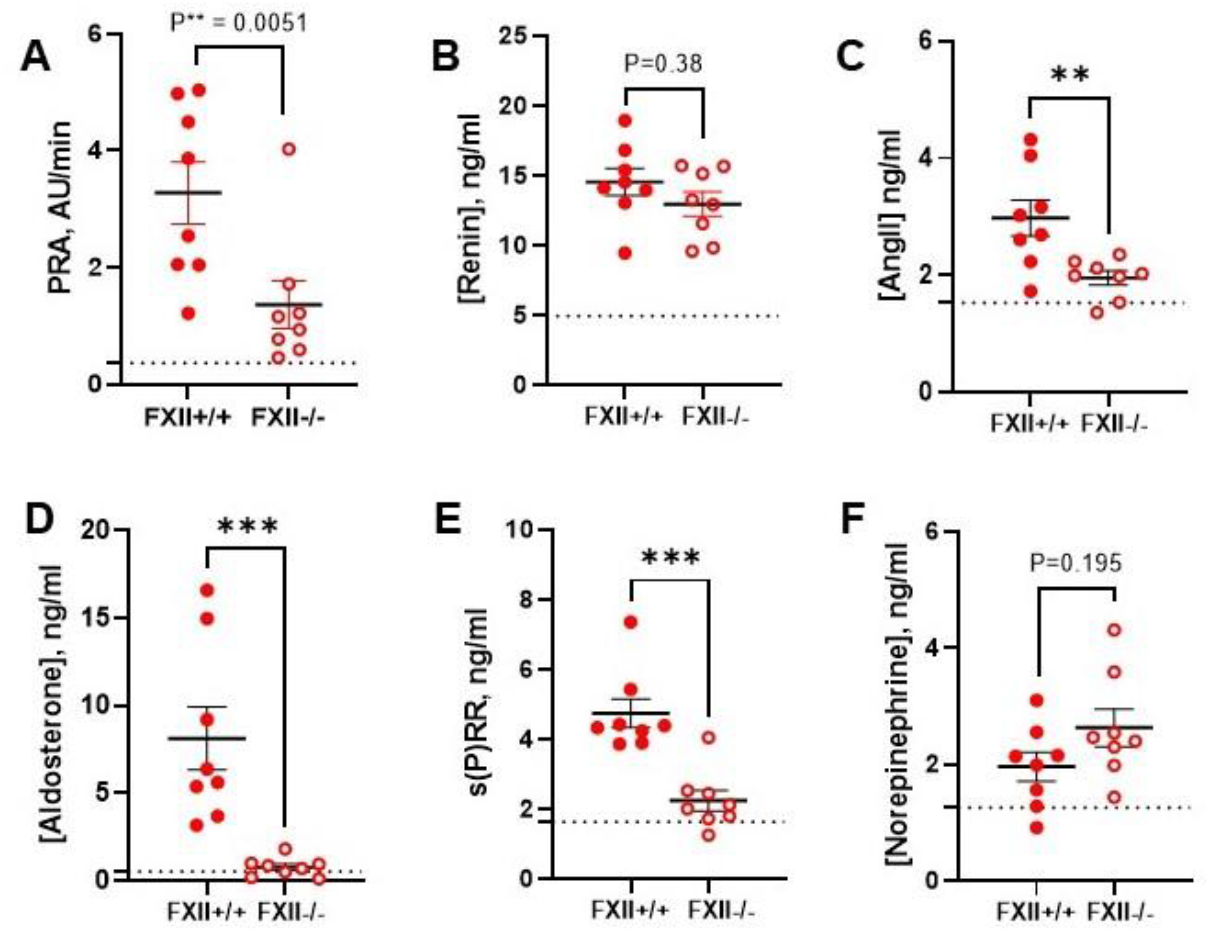
FXII deficiency reduces plasma renin activity and the levels of key members of the renin-mediated cascade of systemic RAAS. In mice with DCM, FXII deficiency (FXII-/-) reduces plasma renin activity, or PRA (**A**) and does not modulate plasma levels of total immunoreactive renin (i.e., renin zymogen) (**B**) in female mice with DCM versus the DCM control group, FXII+/+ (n = 8 per group). FXII deficiency reduces systemic RASS activity: plasma levels of angiotensin II, Ang II (**C**), aldosterone (**D**) and s(P)RR (**E**), while does not modulate norepinephrine (**F**) plasma levels. Studies were performed in a DCM mouse model that is normotensive, without renal dysfunction, highly reproducible and replicates PRA-related observational studies in human DCM-HF. Our data demonstrated that renin activation and sHF progression are enhanced in DCM female mice versus males.^9^ Renin plasma enzymatic activity and plasma biomarkers levels were analyzed as we previously reported.^4,7,9,11,14^ Data were analyzed with unpaired Mann-Whitney test using GraphPad Prism 9.5 software and presented as the mean ± SE. The dotted line shows the levels in wild-type littermates (n = 8) for reference. **P<0.01, ***P<0.001.

Active renin catalyzes the first and rate-limiting step of systemic RAAS activation resulting in angiotensin II production, secretion of aldosterone into circulation, and alterations in the levels of soluble (pro)renin receptor, s(P)RR.^4,7,9^ The reduction of renin activity levels in circulation by FXII-deficiency decreased the activity of systemic RAAS, causing a reduction in circulating angiotensin II, aldosterone, and s(P)RR levels (**Figure 4 C-E**). FXIIa deficiency did not modulate norepinephrine levels in DCM groups (**Figure 4 F**), suggesting that its modulation of plasma renin activity does not affect this process.

The reduced RAAS activation in DCM mice with FXII-deficiency was associated with significantly improved median survival (**Figure 5**), and it delayed onset of sHF (**Figure 6**). In DCM mice, FXII-deficiency significantly improved contractility and pathological heart enlargement, demonstrated by increased ejection fraction (EF), stroke volume, and cardiac output (**Figure 6 A-C**) and decreased in left ventricular mass and heart-to-body weight ratio (**Figure 6 D-E**). It also delayed the timing of sHF onset as it attenuated edema, demonstrated by reduced lung-to-body weight ratio, pleural effusion, and plasma levels of BNP and NT-proANP (**Figure 6 F-I**).

**Figure 5.**
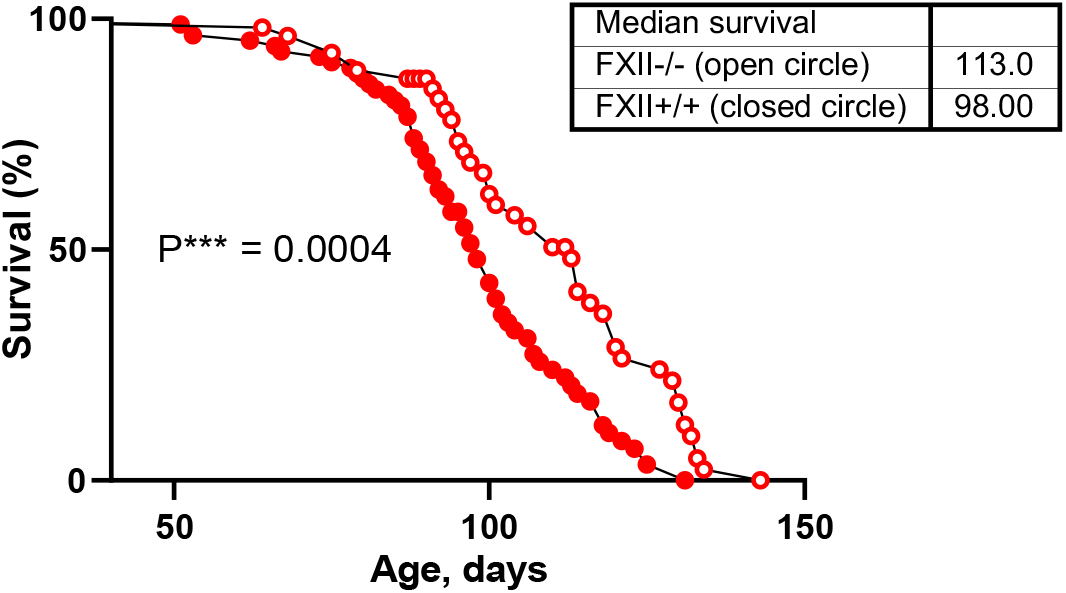
FXII deficiency prolongs life in mice with DCM. FXII deficiency improves the median survival of DCM mice (red open circle) versus the FXII+/+ DCM group (red closed circle). Data was analyzed using Kaplan-Meir survival analysis with the Mantel-Cox test using GraphPad Prism 9.5 software. ***P<0.001.

**Figure 6.**
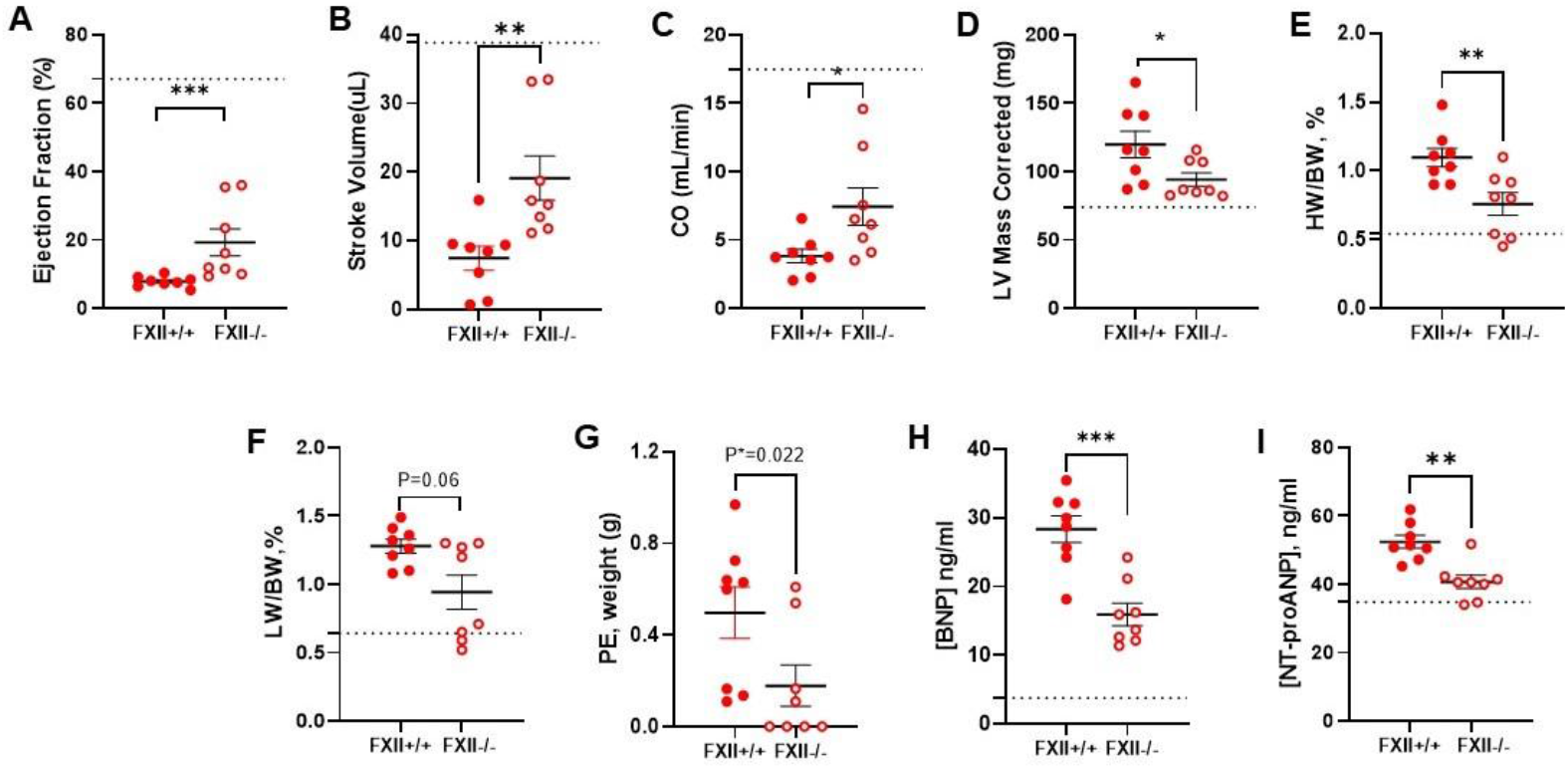
FXII-deficiency attenuates symptomatic HF in mice with DCM. FXII-deficiency prevents progressive contractile dysfunction and pathological heart enlargement in DCM mice versus the DCM control group, FXII+/+ (n = 8 per group): ejection fraction, EF (**A**), stroke volume (**B**), cardiac output, CO (**C**), left-ventricular mass corrected, LV Mass corrected (**D**) and heart weight-to-body weight ratio, HW/BW (**E**). FXII deficiency prevents the development of symptomatic HF defined by edema assessed by difference in lung weight-to-body weight ratio, LW/BW (**F**), pulmonary effusion, PE (**G**) and levels of HF-related plasma biomarkers BNP (**H**) and NT-proANP (**I**). In WT mice, FXII deficiency did not affect survival, systolic function, and remodeling, the plasma levels of HF-biomarkers (BNP and NT-proANP). Data analyzed with unpaired Mann-Whitney test and presented as the mean ± SE. The dotted line shows the levels in wild-type littermates (n = 8) for reference. * P < 0.05, **P<0.01, ***P<0.001.

Due to the prominent role of RAAS in HF, RAAS blockade is one of the cornerstones of HF patients’ guidelines-directed therapies with survival benefits.^3,19^ Our study directly links FXII to renin activity levels in circulation, activation of the systemic RAAS, development of edema, cardiac dilation, and mortality in experimental DCM. Levels of renin activity are reduced by ∼2/3 in FXII-deficient mice with DCM (**Figure 4 A**), indicating that activated FXII is responsible for the majority, but not all renin activation in experimental HF. However, it is unknown whether FXII is required for renin activity generation in other cardiovascular pathologies or physiological conditions. As FXII does not modulate normal hemostasis, targeting FXII’s biological activities is a therapeutic strategy for preventing thrombosis and the inflammatory response in various disease states. The discovery of this novel role of FXII advances our understanding of the regulatory role of the upstream proteolytic network in generating renin activity and regulating systemic RAAS activity during the pathogenesis of HF. Because renin is a master regulator of RAAS, further insights into the mechanism of systemic renin activation and blocking its generation upstream may provide a new therapeutic strategy to control pathologic increases in RAAS by blocking the upstream generation of active renin to prevent HF progression, disability, symptoms, and death.

## Acknowledgments

All animal studies were approved by the University of Arizona’s Institutional Animal Care and Use Committee and performed in accordance with ARRIVE guidelines (Animal Research: Reporting of in Vivo Experiments). The data, analytical methods, and materials that support the findings of this study will be available to other researchers from the corresponding authors on reasonable request.

